# TOM: Transformer Neural Network Optimization of mRNA initiator region

**DOI:** 10.1101/2025.10.30.685412

**Authors:** Ritvik Gupta

**Affiliations:** Independent

## Abstract

Recombinant Protein Production enables scientists to insert custom DNA into host cells to produce specific proteins. Current industry standard tools to optimize recombinant DNA use the Codon Adaptation Index (CAI), yet doing this changes local mRNA secondary structure and Minimum Free Energy (MFE) at the translation initiation region, creating hairpins that block ribosome loading and initiation, thus limiting protein production. mRNA secondary structure around the start codon (disrupting ribosome docking/initiation) is the true rate limiting barrier to protein synthesis, and its 22x more correlated to protein production than CAI. This project introduces TOM, a novel Transformer deep learning model that optimizes mRNA secondary structure at the critical translation initiation region. Over 40 million natural E. Coli initiation sequences were downloaded, where through a rigorous data filtration effort, only 10,000 ideal, non redundant, and naturally occurring sequences were used to train the model. TOM is benchmarked on MFE, adenine count, codon usage, and a negative element analysis against industry standard optimizers and significantly outperforms. TOMs improvement of the MFE at the initiation region offers a significant increase in protein production by addressing the rate-limiting step of translation initiation.

## Background

Custom proteins are synthetically created through cell factories in a process known as recombinant protein production [8]. The application of recombinant proteins is endless. While recombinant proteins are mainly used for therapeutic efforts-like vaccines, anti-cancer/viral peptides, antibodies, and essential hormones [9, 10, 11, 12]-there is promising application of proteins in environmental issues. For example, proteins have been discovered that can biodegrade plastic efficiently [13]. Therefore, it is essential that these life-saving proteins can be efficiently mass produced.

The process of producing custom proteins in host cells generally goes as follows: designing a synthetic plasmid that contains the gene encoding the protein, incorporating the plasmid into host cells, and utilizing the cell’s transcription and translation machinery to produce the target protein [7]. E. coli is a preferred method of expressing proteins because of its cheap but very large scale production [6]. There are many factors that affect the protein expression (production) levels of these synthetic genes in heterologous hosts. Due to this, optimizing plasmid DNA for increased protein expression is a frontier of large potential.

Genetic code is degenerate. 64 codons encode only 20 amino acids, so there are synonymous codons that encode the same amino acid. For example, the codons ACA, ACG, ACC, and ACU all encode the Threonine (Thr) amino acid. Codon optimization is the common approach to increase protein expression. This process chooses synonymous codons that are more frequently used in the host organism, typically based on codon usage bias. Usually, models will optimize the Codon Adaptation Index (CAI) metric to achieve this. By selecting codons that match the host’s highly abundant tRNAs, translation elongation can be improved, potentially leading to more expression. [14, 15]. There exist many models that optimize CAI in the coding sequence to increase expression [16, 17].

Protein expression happens in three phases: initiation, elongation, and termination [20]. The common approach to optimizing gene expression-CAI optimization-only benefits the translation elongation step, which actually isn’t the rate limiting step in protein synthesis. Translation initiation has been accepted as the rate limiting step for gene expression [18]. Evolutionary analysis has shown that nature doesn’t select for higher CAI if the translation initiation is weak, since increasing elongation efficiency under weak initiation circumstances wouldn’t benefit expression [19]. Under controlled E. coli expression experiments, translation initiation is 30x more correlated to protein expression than CAI [22]. The study concludes that codon usage is likely to be more connected to cellular fitness, and translation initiation is the key determining factor of gene expression. This highlights the importance of optimizing translation initiation before translation elongation in order to increase protein expression.

Translation initiation is critical in protein synthesis. It involves the assembly of the ribosomal subunits, mRNA, and initiator tRNA, which is a complex process regulated by various initiation factors [1, 2]. Translation initiation is the rate-limiting step in protein synthesis, and therefore has the largest impact on protein production [3]. mRNA secondary structure minimum free energy (MFE) of the region surrounding the initiator codon has the most significant impact on translation initiation [3]. Secondary structures in the 5’ upstream untranslated region (UTR) and the Coding Sequence (CDS) region downstream the start codon, significantly impacts ribosome access. Stable secondary structures (e.g., hairpins with high GC content) create energy barriers that impede ribosome binding and scanning. In contrast, weaker secondary structures (lower Minimum Free Energy, MFE) allow ribosomes to access the initiator mRNA more efficiently. This is because the ribosome must unwind these structures during initiation, and higher stability requires more energy and time, slowing down the process [4]. This relationship between MFE and translation initiation and overall expression has driven evolutionary selection towards weaker 5’ mRNA structure for efficient translation initiation, which demonstrates the importance of unstable mRNA structures for increased translation initiation rates [5]. In E. coli, translation initiation is highly correlated to the total protein yield (r^2>.98) [3].

Therefore, to increase recombinant protein expression, there should be a focus to change the target mRNA sequence to be optimized for translation initiation.

There are very few models that focus on optimizing for translation initiation in order to increase gene expression. The industry standard is RBS Designer, which creates the 5’ untranslated region [23]. However, the model does not account for the CDS just downstream the initiator codon, which plays a major role in translation initiation [24]. Because multiple codons encode amino acids, selectively choosing synonymous codons that encode the same amino acid but could have different effects on MFE would consequently influence translation initiation. Therefore, there is opportunity to co-optimize both the UTR and CDS surrounding the initiator codon to increase expression.

Experimental evidence has shown that the MFE of the −4 to +39 sequence relative to the start codon has the most statistically significant impact on gene expression [22]. Additionally, in the region close to the translation initiation site, mRNA secondary structure, not CAI, is the key determinant of protein production [25]. This highlights the flaws of only optimizing CAI in the entire CDS, as this ignores the importance of the initial few codons’ relation to translation initiation. It also shows that using ‘rare’ codons (low CAI) in earlier parts of the CDS shouldn’t harm protein expression if the rare codon contributes to better MFE in the translation initiation region.

Understanding the evolutionary context of 5’ UTR sequences and codon usage near the start codon could help create translation initiation regions that are more similar to E. coli native gene initiatory sequences. This is crucial because recombinant protein production in heterologous hosts can sometimes lead to cellular stress or toxicity if the introduced sequences deviate significantly from naturally evolved sequences [16]. E. coli has adapted its translation initiation mechanisms to efficiently process its native genes, and disrupting this balance with foreign or highly engineered sequences can lead to unintended consequences.

To capture the underlying genetic context of E. coli during translation initiation, deep learning models like Transformers can be leveraged. Transformers have revolutionized computational biology by significantly enhancing the understanding of sequential dependencies in genetic data [27]. In this research project, a Transformer-based deep learning model was trained on native E. coli translation initiation sequences, specifically filtered for optimal minimum free energy (MFE) conditions. By learning the sequential patterns associated with efficient translation initiation within E. coli, the model can generalize this knowledge to generate novel mRNA initiation sequences optimized for high expression.

## Methodology

### Data Preprocessing

I utilized the National Center for Biotechnology Information’s (NCBI) command line tool to download all E. coli genomes that have been sequences. I downloaded the standard genome file, but also the cds_from_genomic file which contained the coding sequences for proteins within the genome. To isolate only the −4 to +39 sequence relative to the start codon for each CDS, I parsed through each CDS in the cds_from_genomic file, and found the sequence within the genome file. I then found the 4 nucleotides behind the CDS within the genome file. These sequences were saved as a .csv file. The csv file contained 42,333,113 sequences. I then filtered through these sequences to get rid of “Psuedo=True” sequences as these genes don’t encode the expected translation, which thus don’t represent the data required for the model [28]. This reduced the amount of sequences to 40,950,651. Finally, CDS that didn’t have the initiator codon as either ATG, TTG, or GTG were removed because of their specialized nature and their inefficiency at translation [29]. Now the sequence count was 40,752,883.

If AI is trained on a redundant dataset, it becomes too specialized and fails to generalize for real-world scenarios [30]. To address this, I clustered the nucleotide sequences with 90% similarity using the MMseq2 linclust tool [31]. For the clustering process, I only based it on the coding regions. I deliberately excluded the −4 regulatory region from clustering, as sequences with optimized MFE may share similar regulatory elements. Removing them could inadvertently discard valuable training examples. Additionally, since the model takes protein sequences as input and generates DNA sequences as output, clustering the coding DNA sequences ensures a diverse training set relative to the proteins, allowing the model to learn from a broad range of potential protein inputs. Using only the representative sequences of the clusters, the data had reduced down to 138,123 sequences. From this, mRNA folding energies were calculated using the ViennaRNA package [36]. Finally, data was selected for ideal MFE conditions. This was done by selecting the 10,000 highest MFE sequences. This process created a non-redundant, representative, and ideal training dataset.

**Figure 1.**
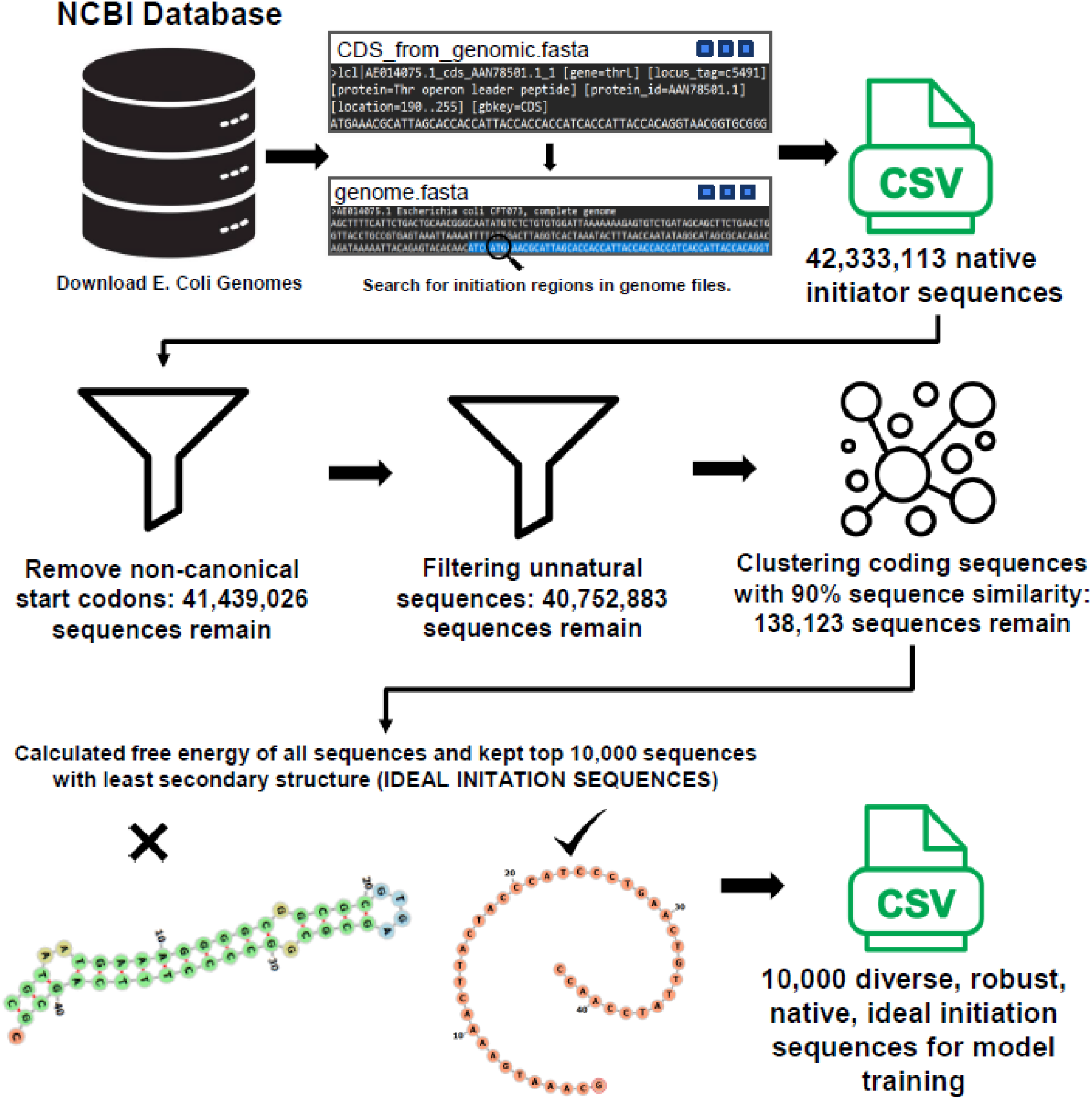

### Model Training

The development and training of the transformer-based model for protein-to-mRNA sequence prediction were conducted using the Torch [32] deep learning framework. This section outlines the data preprocessing steps, model architecture, and training procedures that were undertaken to build an effective sequence translation system.

#### Data Preprocessing

The dataset used in this study consisted of paired protein sequences and their corresponding mRNA sequences. These sequences were extracted from a curated dataset stored in a CSV file. Given that sequence data is inherently categorical, it was necessary to convert the characters into numerical representations to facilitate processing by the model, which is commonly done in transformer based models in computational biology [27]. Character-level tokenization was applied, where each unique character in both protein and mRNA sequences was assigned a corresponding numerical index.

For protein sequences, a vocabulary was constructed from the distinct amino acid characters present in the dataset. An additional padding token () was introduced to ensure uniform sequence lengths, thereby enabling efficient batch processing. Similarly, for mRNA sequences, a vocabulary was built with unique nucleotide characters, supplemented by special tokens such as start-of-sequence () and end-of-sequence () markers. These markers were introduced to guide the model in learning sequence generation effectively. Each protein sequence was padded to a length of 13, while mRNA sequences were padded to a length of 45 to maintain consistency across input samples.

A transformer-based neural network was implemented to perform sequence-to-sequence translation between protein and mRNA sequences. The model architecture was inspired by the standard transformer framework, which has demonstrated superior performance in various computational biology processing tasks [33, 34]. The network comprised an encoder-decoder structure, where the encoder processed the protein sequence embeddings, and the decoder generated the corresponding mRNA sequences.

The encoder module consisted of an embedding layer followed by multiple transformer encoder layers. The embedding layer transformed input indices into high-dimensional vector representations, while positional encoding was added to incorporate sequential dependencies. The encoded protein sequence representations were subsequently fed into a transformer-based decoder. The decoder architecture mirrored the encoder, comprising embedding layers and transformer decoder blocks, with an additional masked self-attention mechanism to regulate token dependencies during autoregressive sequence generation. A final linear projection layer converts the decoder’s output into logits over the mRNA vocabulary.

A summary of the model architecture can be seen in supplementary file 2.

#### Training Procedure

To train the transformer model, the dataset was split into training and validation sets, with 90% of the data allocated for training and the remaining 10% used for validation. The data was loaded into the PyTorch dataloaders to facilitate efficient batch processing. A batch size of 32 was used for both training and validation.

The model was optimized using the Adam optimizer [35] with a learning rate of 5e-4. Cross-entropy loss, with padding tokens ignored, was employed as the loss function to measure the discrepancy between the predicted and actual mRNA sequences. During training, the decoder utilized teacher forcing by taking the ground truth sequence (shifted by one position) as input to predict the next token in the sequence. Training was conducted over ten epochs, with loss values computed for both training and validation sets. To monitor training progress, loss values were recorded at each epoch, and performance was evaluated based on the reduction in loss over successive iterations. The model exhibited progressive improvement in learning meaningful mappings between protein and mRNA sequences, as reflected by the decreasing loss values. Additionally, validation loss trends were closely monitored to detect potential overfitting.

**Figure 3.**
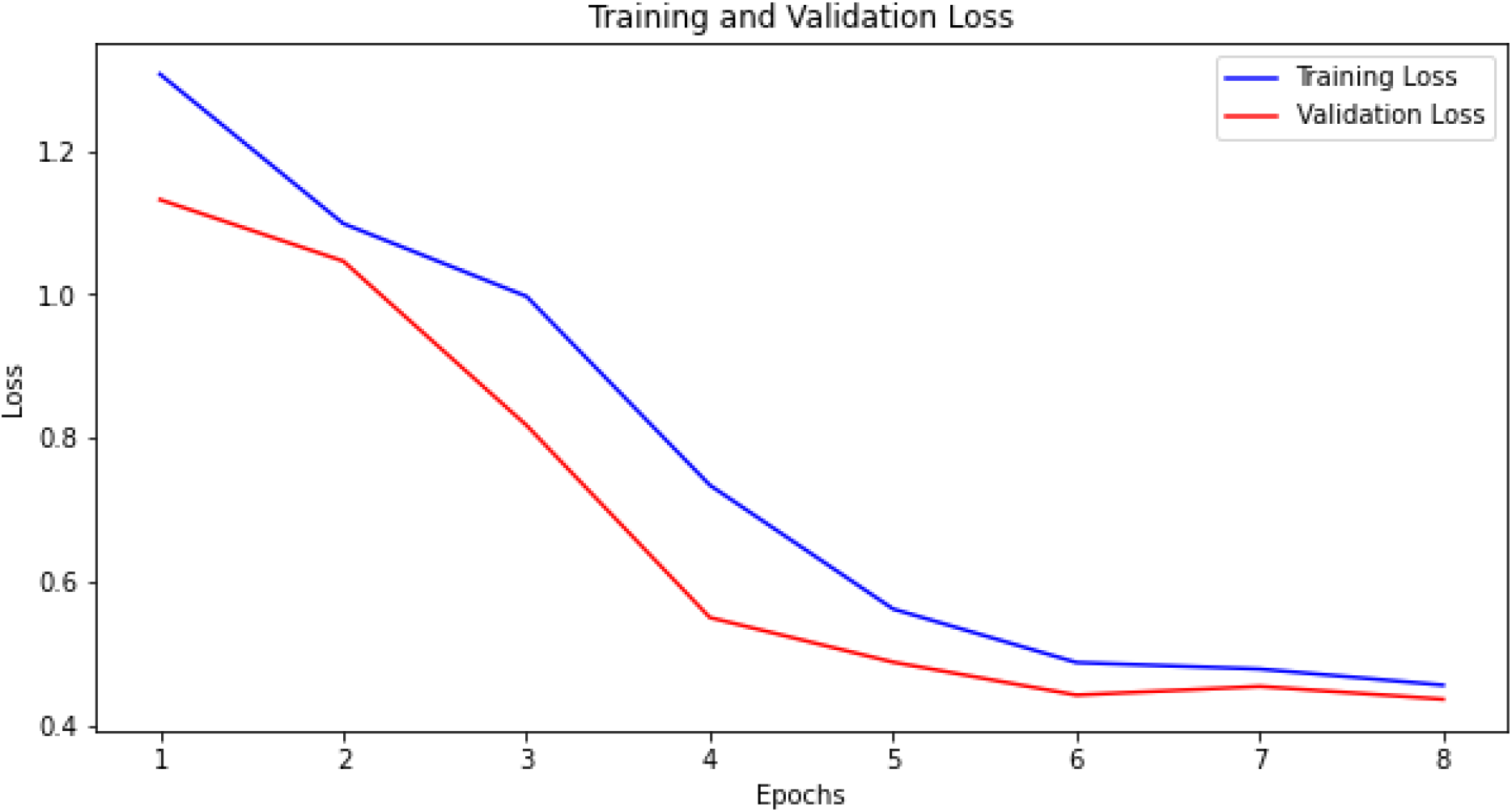
The validation loss was less than the training loss, which means that the model did not overfit on training data and is generalizable on unseen data.

### Results/Model Validation

Given the computational expense of running a large number of sequences through transformer models, the number of sequences was constrained. The computational limitations were taken into account during the validation process, and efforts were made to ensure the results were robust despite these constraints.

To assess the robustness of the model I randomly selected 500 mRNA sequences from the clustered representative dataset that had an MFE that was lower than the ideal threshold (MFE > −5). This choice was based on the observation that the majority of sequences in the clustered representatives dataset fell within this range.

To statistically validate whether the model achieved significant improvements in MFE in the significant region for initiation (−4 to +37) through optimized UTR/codon choices, I converted the test mRNA sequences into proteins and ran them through TOM to obtain the optimized mRNA sequences. I performed two different statistical tests. Paired t-test results and Wilcoxon signed-rank.

**Figure 4.**
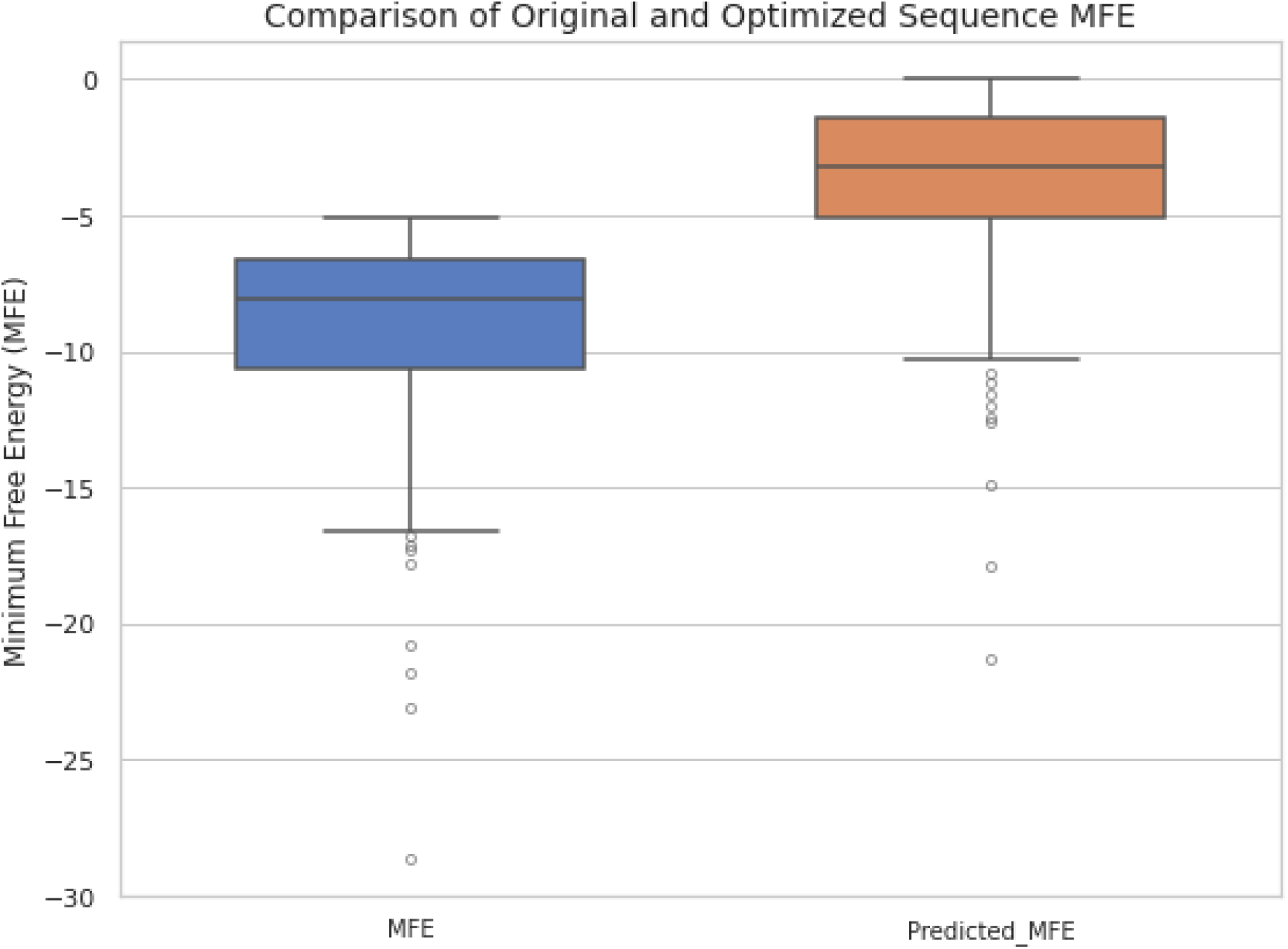
Paired t-test results: t-statistic = −32.45694813127643 p-value = 4.5020536104376186e-125 Wilcoxon signed-rank test results: Statistic = 2414.5 p-value = 1.9170542472302483e-77

**When extrapolating the average MFE increase by TOM (∼5.53 kcal/mol) to expression studies that isolated all variables except mRNA secondary structure MFE at the −4 to +39 region, protein production is estimated to improve by over 3-fold [22]**

Based on my thorough literature review, no existing models besides TOM are capable of optimizing initiator coding sequence (CDS) MFE along with the UTR to enhance translation initiation. Researchers typically optimize the entire CDS for codon adaptation index (CAI) using industry-standard tools. However, doing this fails to consider the critical role of the initiator CDS in translation initiation. The CAI of the initiator sequence has a much weaker correlation with gene expression compared to its minimum free energy (MFE), as MFE of this region directly influences initiation efficiency and overall expression levels [24,25]. This limitation made direct comparisons challenging, as no models currently exist that specifically optimize translation initiation via CDS design.

To evaluate whether my model enhances translation initiation, I optimized the MFE of the initiator CDS using TOM and benchmarked its performance against industry-standard models designed to increase protein expression. Since these models focus solely on CAI and do not account for untranslated regions (UTRs), I compared only the CDS optimization results, excluding UTR-related effects. Notably, research indicates that the MFE of the first few codons—independent of the UTR—is also strongly associated with expression levels [25]. By increasing MFE in this critical region, TOM provides a novel approach to using synonymous codons to improve translation initiation, distinguishing it from the current state of the art optimization methods.

**Figure 5:**
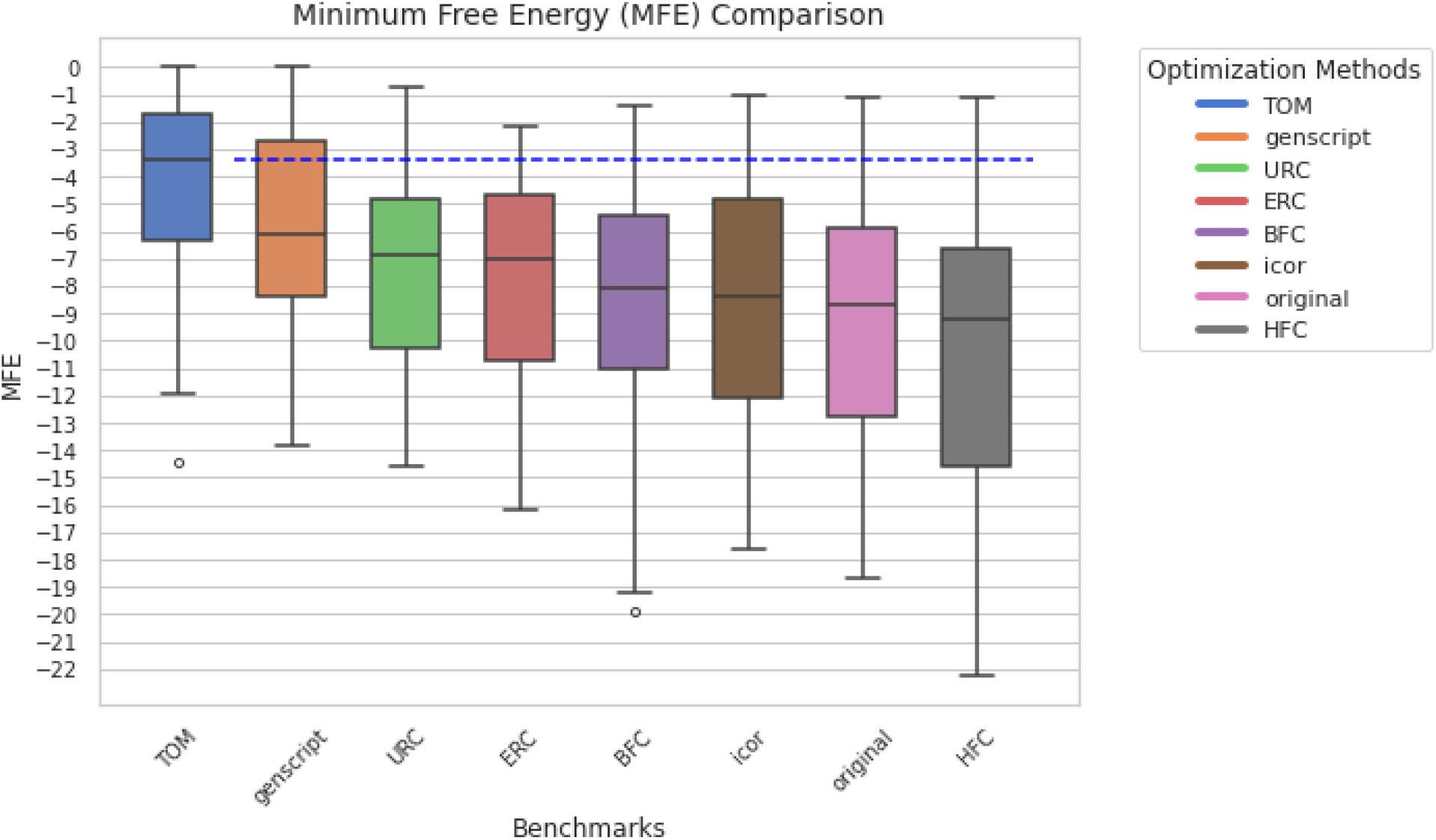
Comparison of TOM with different protein expression optimizers (n=37). The dataset for the optimized sequences by the other models was taken from [16]. TOM significantly increases MFE of the initiator CDS (p<.001) compared to the genscript CDS optimizer based on a paired t test.

High Adenine count has been naturally favored in the first few codons, and it generally improves protein production [44]

**Figure 6.**
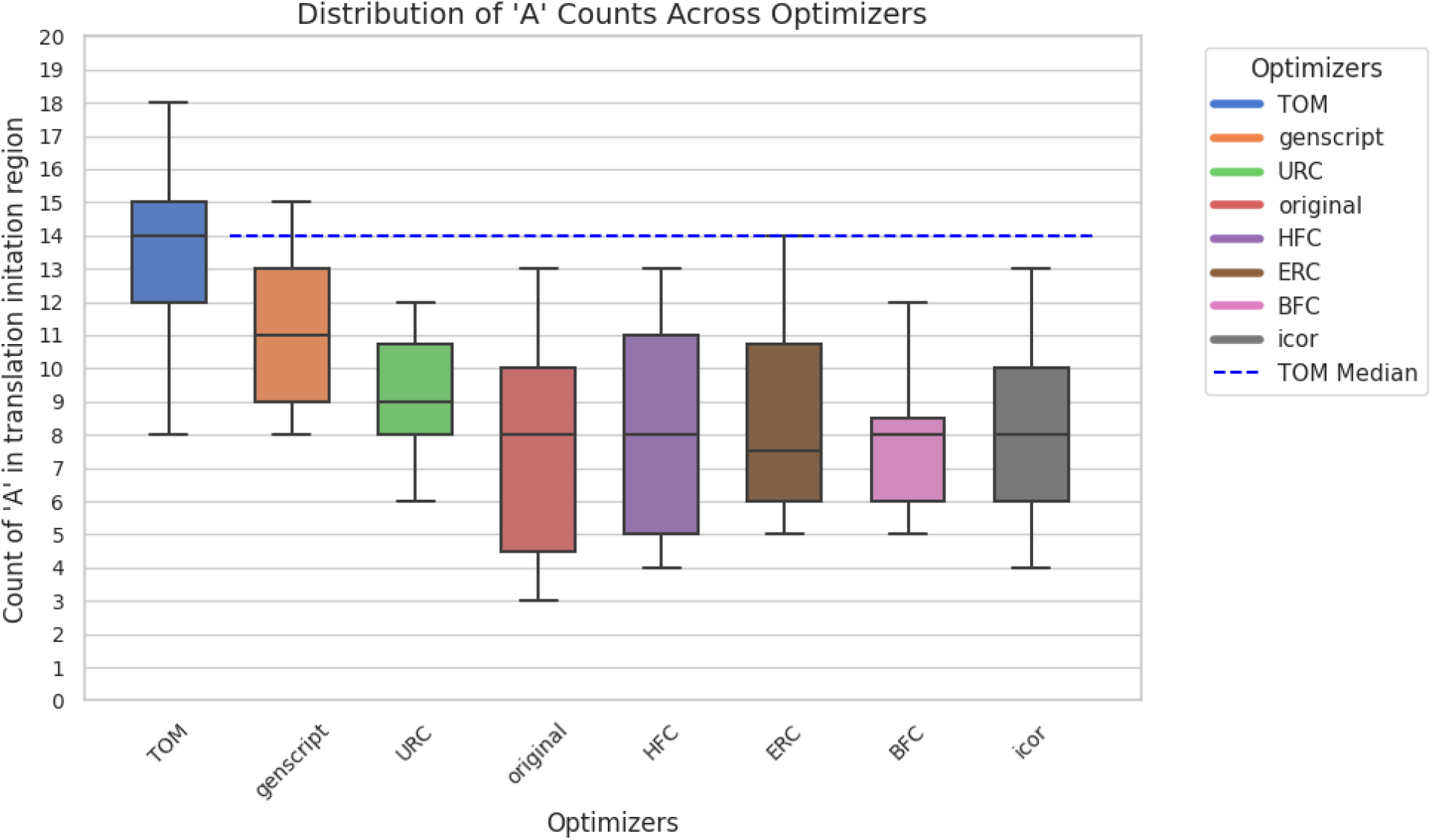
Comparison of TOM with different protein expression optimizers (n=37). The dataset for the optimized sequences by the other models was taken from [16]. TOM significantly increases Adenine counts of the initiator CDS (p<.001) compared to the genscript CDS optimizer based on a paired t test.

I wanted to see if optimizing the initiation region with TOM and optimizing the elongation region with optimizers designed to improve CAI was feasible. To do this, I analyzed if incorporating TOM optimization for the initiation region could be implemented into a full sequence optimized by some other optimizer without introducing negative elements.

**Figure 8.**
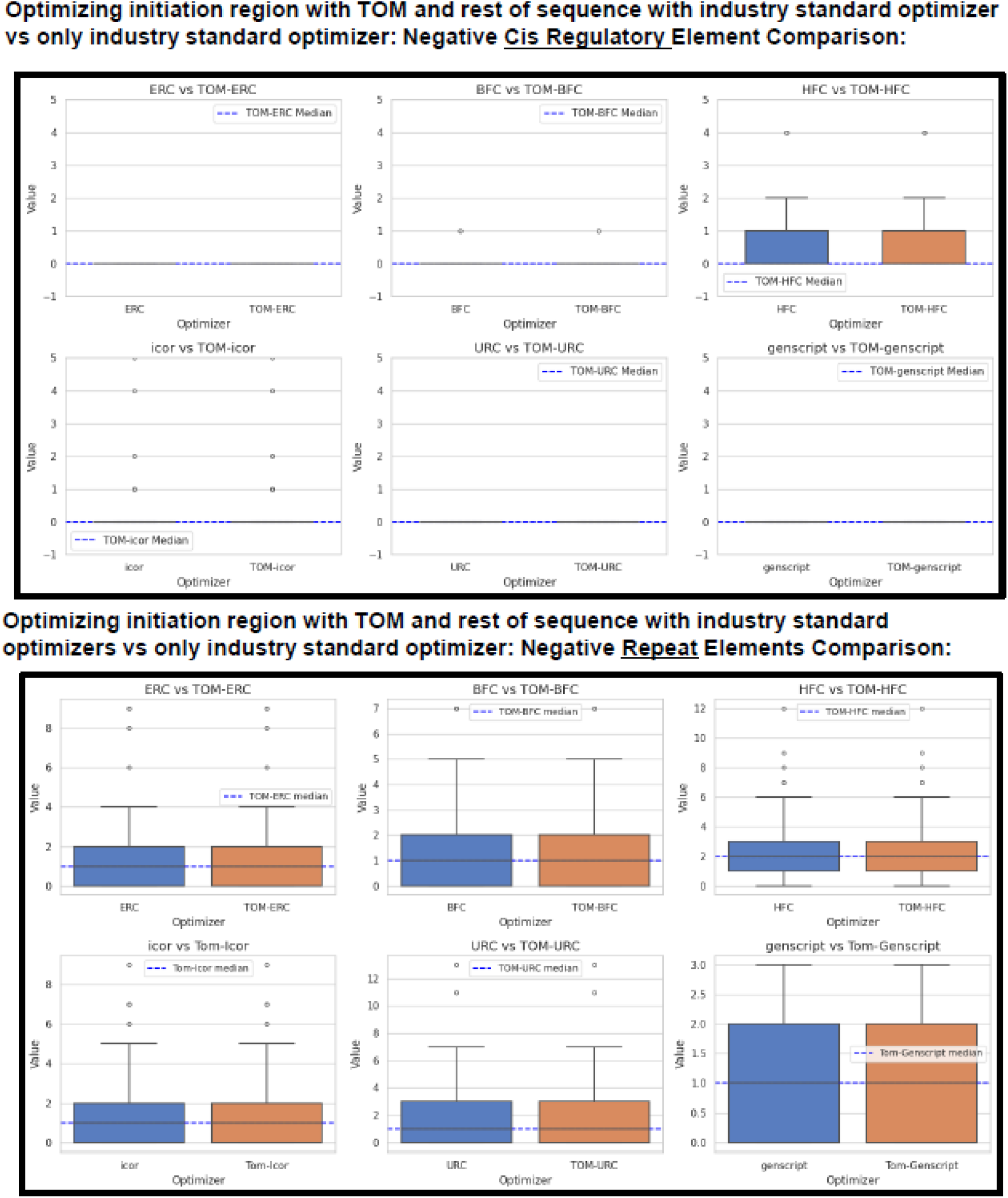
Comparison of negative element count when sequence is co-optimized with TOM/ without TOM. TOM does not introduce any new negative elements.

### Mutational Analysis

If even one non-synonymous codon was predicted by the model, that would’ve caused the encoded protein sequence to be different. In both validation tests, there was a 0.00% mutational rate that would’ve otherwise caused a change in the encoded protein sequence. This demonstrates that the model clearly understands the relationship between mRNA and amino acid sequences.

### Example Usage of TOM

Insulin is a hormone protein that is recombinantly produced as a therapeutic to diabetes [12]. To exemplify the application of TOM, here is an example usage of TOM on the therapeutic insulin translation initiation sequence

**Figure 9.**
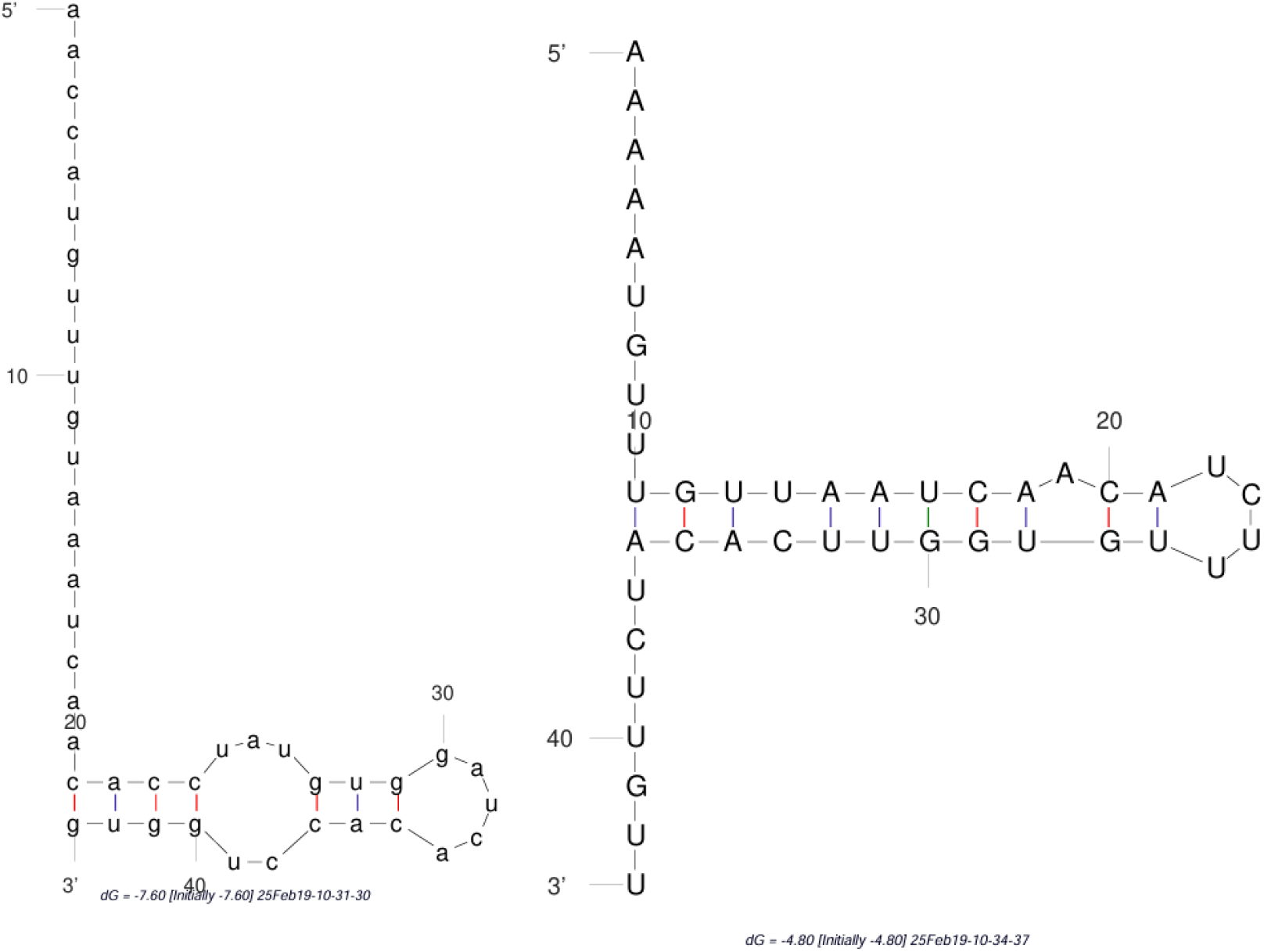
Comparison of codon optimized industry standard insulin initiation sequence (left) with TOM optimization of insulin initiation sequence (right). The folding was done via mFold [40]

The codon optimized sequence of insulin has a folding energy of −7.6 kcal/mol. This sequence is what is usually used in recombinant expression systems. After optimizing with TOM, the sequence has a folding energy of −4.6 kcal/mol. When extrapolating the MFE increase by TOM (3 kcal/mol) to expression studies that isolated all variables except mRNA secondary structure MFE at the −4 to +39 region, insulin production is estimated to improve [22] with TOM initiation optimization. Considering that 1 million Americans ration insulin, leading to ketoacidosis and thus more hospitalizations [38, 39], increasing insulin production could help mitigate the issues of high-cost pharmaceuticals.

### Environmental Application of TOM

Since the 1960s, approximately 6,300 million metric tons (Mt) of plastic waste have been produced, with a mere 9% being recycled, while the rest accumulates in landfills or the natural environment. If current trends persist, projections suggest that around 12,000 Mt of plastic will be landfilled by 2050, exacerbating environmental degradation. This growing plastic waste poses severe threats to ecosystems, wildlife, and human health, as plastics take hundreds of years to break down. PETase, a plastic-degrading enzyme, offers a promising solution to this crisis. PETase is recombinantly produced [41].

Test example of TOM vs Genscript: Ribosomal protein S1 (mRNA binder protein of ribosome) docked to mRNA translation initiator region encoding Plastic Digesting Enzyme PETase:

**Figure 10.**
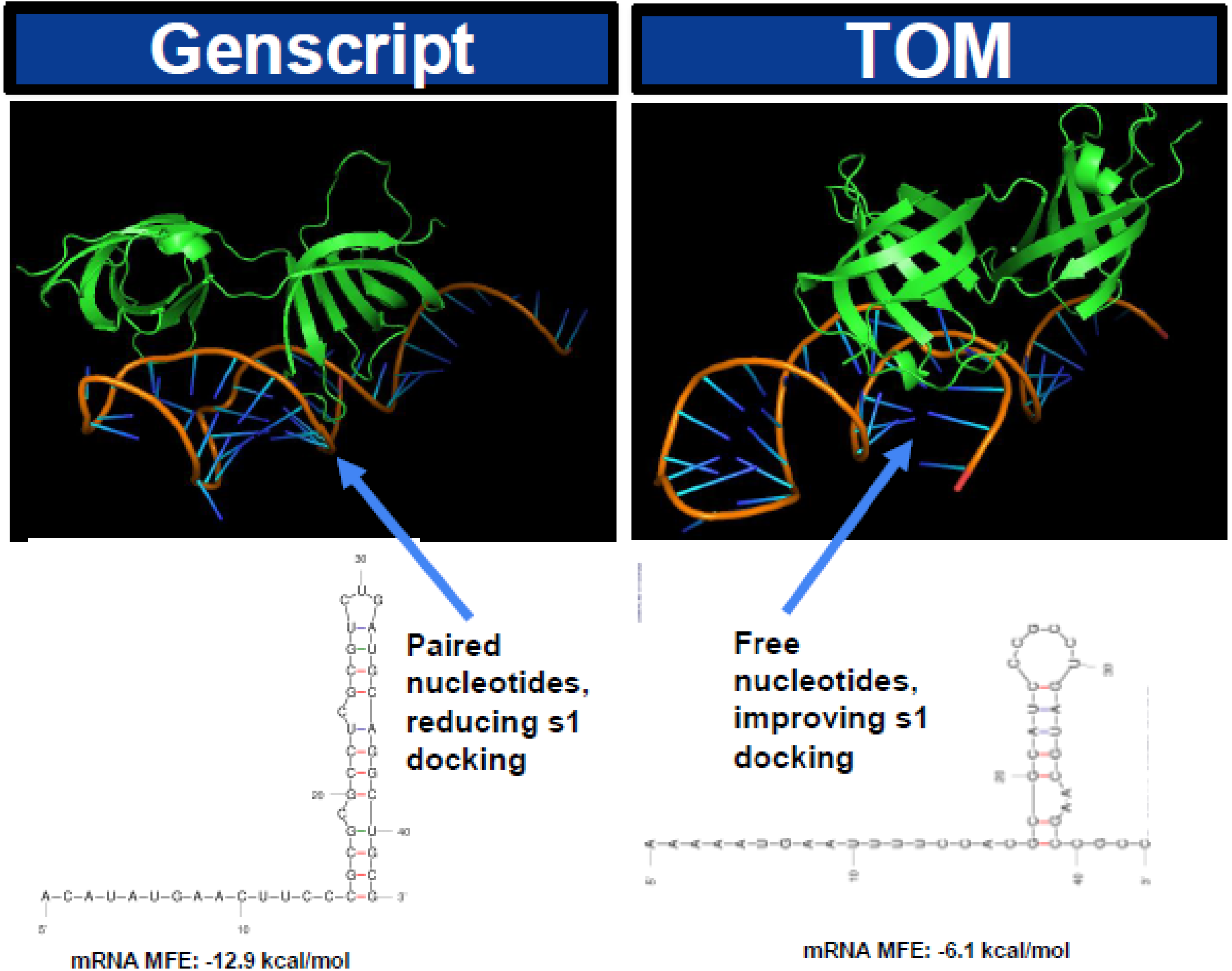
TOM improves initiator mRNA binding to ribosomal protein by ∼5 kcal/mol. TOM was able to choose synonymous codons that reduced mRNA secondary structure, which freed up nucleotides. This suggests that TOM improves ribosomal accessibility to mRNA, thus improving initiation. These are just two examples. TOM can be applied to any other protein to increase its production.

## Conclusions/Impact

This research highlights the significance of optimizing translation initiation, rather than solely focusing on CAI, to enhance recombinant protein expression in E. coli. By leveraging deep learning, this study captures the underlying genetic context of ideal native E. coli translation initiation sequences to generate optimized mRNA designs of the initiation region. The model’s ability to co-optimize both the 5’ UTR and the coding region downstream of the start codon provides a novel approach that surpasses the current state of the art methods. Because of the fact that TOM does not introduce any negative elements into sequences, it can be used to optimize only the initiation region, while the rest of the sequence would be optimized by another optimizer. Given that translation initiation is the rate-limiting step and highly correlated with overall protein yield, this approach offers a more biologically relevant and effective strategy for increasing expression levels. This research could be further extended to optimize translation initiation across different expression hosts and even mRNA based vaccines.

The ability to significantly enhance protein expression has profound implications for therapeutics. For instance, optimized protein expression can streamline the production of therapeutic proteins, such as monoclonal antibodies, cytokines, and enzymes, reducing costs and improving scalability for treatments targeting cancer, autoimmune diseases, and rare genetic disorders. In vaccine development, enhanced expression of antigenic proteins can accelerate the production of vaccine candidates.

The immediate next step is experimental validation of the generated sequences to confirm their efficacy in enhancing protein expression. Beyond this, the research can be extended to optimize translation initiation across diverse expression hosts, including mammalian and yeast systems, further advancing the field of synthetic biology and recombinant protein production. By addressing the bottleneck of translation initiation, this research not only improves protein yields but also enhances the efficiency, affordability, and accessibility of biotherapeutics.

## Acknowledgments

This research used publicly available genomic datasets and did not involve human participants, animals, or clinical trials. No ethical approval was required.

